# Hyperexcitability in the olfactory bulb and impaired fine odor discrimination in the *Fmr1* KO mouse model of fragile X syndrome

**DOI:** 10.1101/2023.04.10.536251

**Authors:** Praveen Kuruppath, Lin Xue, Frederic Pouille, Shelly T. Jones, Nathan E. Schoppa

**Author notes:** <u>E-mail of corresponding author</u>. <u>Conflict of Interest</u>: None of the authors have an actual or potential conflict of interest in relation to this manuscript.

## Abstract

Fragile X syndrome (FXS) is the single most common monogenetic cause of autism spectrum disorders in humans. FXS is caused by loss of expression of the Fragile X mental retardation protein (FMRP), an mRNA-binding protein encoded on the X chromosome involved in suppressing protein translation. Sensory processing deficits have been a major focus of studies of FXS in both humans and rodent models of FXS, but olfactory deficits remain poorly understood. Here we conducted experiments in wild-type and *Fmr1* KO (*Fmr1^-/y^*) mice (males) that lack expression of the gene encoding FMRP to assess olfactory circuit and behavioral abnormalities. In patch-clamp recordings conducted in slices of the olfactory bulb, output mitral cells (MCs) in *Fmr1* KO mice displayed greatly enhanced excitation, as evidenced by a much higher rate of occurrence of spontaneous network-level events known as long-lasting depolarizations (LLDs). The higher probability of LLDs did not appear to reflect changes in inhibitory connections onto MCs but rather enhanced spontaneous excitation of external tufted cells (eTCs) that provide feedforward excitation onto MCs within glomeruli. In addition, in a go/no-go operant discrimination paradigm, we found that *Fmr1* KO mice displayed impaired discrimination of odors in difficult tasks that involved odor mixtures but not altered discrimination of monomolecular odors. We suggest that the higher excitability of MCs in *Fmr1* KO mice may impair fine odor discrimination by broadening odor tuning curves of MCs and/or altering synchronized oscillations through changes in transient inhibition.

**Significance Statement:** Fragile X syndrome (FXS) in humans is associated with a range of debilitating deficits including aberrant sensory processing. One sensory system that has received comparatively little attention in studies in animal models of FXS is olfaction. Here, we report the first comprehensive physiological analysis of circuit defects in the olfactory bulb in the commonly-used *Fmr1* knockout (KO) mouse model of FXS. Our studies indicate that *Fmr1* KO alters the local excitation/inhibition balance in the bulb – similar to what *Fmr1* KO does in other brain circuits – but through a novel mechanism that involves enhanced feedforward excitatory drive. Furthermore, *Fmr1* KO mice display behavioral impairments in fine odor discrimination, an effect that may be explained by enhanced neural excitability.

## Introduction

Fragile X syndrome (FXS) is the single most common monogenetic cause of autism spectrum disorders (ASDs) in humans. Individuals with FXS can exhibit a range of debilitating deficits in cognitive abilities, social interactions and communication, hyperactivity and anxiety, and sensory processing (Hagerman et al., 2017; Mila et al., 2018). Deficits in sensory processing, often hypersensitivity to stimuli, can contribute to social interaction deficits. FXS results from loss of expression of the Fragile X mental retardation protein (FMRP), an mRNA-binding protein encoded on the X chromosome involved in suppressing protein translation. Previous studies in FXS patients and animal models of FXS have demonstrated that impaired expression of FMRP causes defects in neuronal branching and dendritic spine morphology (Davis and Broadie, 2017; Bagni and Zukin, 2019). Some of the most marked functional changes are within circuits involved in sensory processing (Sinclair et al., 2017; Rais et al., 2018). For example, in the visual, auditory, and tactile systems, *Fmr1* KO mice that lack expression of the gene encoding FMRP display enhanced neural responsiveness, widening of receptive fields, and/or behavioral hypersensitivity (Chen and Toth, 2001; Arnett et al., 2014; Rotschafer and Razak, 2014; Juczewski et al., 2016; Wen et al., 2019; Razak et al., 2021; Yang et al., 2022).

One sensory system that has received comparatively less attention in rodent models of FXS is olfaction (reviewed in (Bodaleo et al., 2019)). Only a few publications have reported olfactory behavioral deficits in *Fmr1* KO mice, with somewhat conflicting results as to whether KO alters olfactory sensitivity or discrimination (Larson et al., 2008; Schilit Nitenson et al., 2015). Similarly, there are only a few reports of effects of *Fmr1* KO on olfactory neural circuits in rodents. For example, Galvez and co-workers (Galvez et al., 2005) reported that *Fmr1* KO can cause a modest increase in the number of apical dendrites of output mitral cells (MCs) of the olfactory bulb (in at least one mouse strain). Also, KO can cause moderate increases in the density and length of spines in adult-born GABAergic granule cells (Scotto-Lomassese et al., 2011). Whether these structural changes manifest in physiological changes in bulb output neurons is unclear, and *Fmr1* KO effects on most neurons in the bulb are completely unstudied. The relative paucity of studies of olfaction in *Fmr1* KO mice is surprising. FMRP is highly expressed throughout the bulb (Hinds et al., 1993; Christie et al., 2009; Akins et al., 2012; Brackett et al., 2013; Korsak et al., 2017), including within Fragile X granules that are implicated in the transport of mRNA to synaptic sites (Lai et al., 2020). In addition, olfaction is expected to be an excellent model system to study FXS deficits in general. Olfaction is a critical sensory system for mice, playing essential roles both in finding food and reproduction (Bhattacharyya and Bhalla, 2015; Gire et al., 2016; Ishii and Touhara, 2019), and, as such, behavioral experiments that assess olfactory function are well-developed and robust (Bodyak and Slotnick, 1999). Also, at least for the broad class of ASDs, there is increasing evidence that humans suffer olfactory deficits (Dudova et al., 2011; Lane et al., 2014; Rozenkrantz et al., 2015; Boudjarane et al., 2017; Koehler et al., 2018).

Here, we sought to examine olfactory dysfunction in *Fmr1* KO mice using both physiological and behavioral approaches. In our electrophysiological analyses, conducted in olfactory bulb slices, we found that *Frm1* KO caused a drastic increase in spontaneous excitatory events in MCs that reflect activity in a glomerular network. This increase was due at least in part to enhanced activity of external tufted cells that provide feedforward excitation onto MCs (De Saint Jan et al., 2009; Najac et al., 2011; Gire et al., 2012). In behavioral studies involving a go/no-go operant discrimination task, we found that *Fmr1* KO impaired the ability of mice to discriminate monomolecular odors from odor mixtures, but did not impair easier discrimination tasks.

## Materials and Methods

### Ethical approval and experimental animals

All experiments were approved by the Institutional Animal Care and Use Committee at the University of Colorado Anschutz Medical Campus in accordance with guidelines set by the U.S. Department of Health and Human Services and outlined in the Public Health Service Policy on Humane Care and Use of Laboratory Animals.

*Fmr1* KO mice (C57BL/6 background) were acquired from Jackson Laboratories (Bar Harbor, ME, USA –JAX stock number 003025) and bred in-house. Experimental *Fmr1* KO mice (hemizygous *Fmr1^–/y^* males) were generated by crossing homozygous *Fmr1* KO (*Fmr1^–/–^*) females with hemizygous *Fmr1^–/y^* males. Wild-type mice (C57BL/6 males) were obtained from Charles River Labs (Wilmington, MA). Mice were maintained on a 12 h light/dark cycle with food and water ad libitum except under training and testing conditions (see below). For slice experiments, mice were at P21-P42. For behavioral studies, mice were approximately 6 weeks old at start of experiments.

### Olfactory bulb slice preparation and electrophysiological recordings

Acute horizontal olfactory bulb slices (300-350 μm thickness) were prepared from mice following isoflurane anesthesia and decapitation as described previously (Zak and Schoppa, 2022). Experiments were carried out under an upright Zeiss Axioskop 2 FS Plus microscope (Carl Zeiss MicroImaging) fitted with differential interference contrast (DIC) video microscopy and a CCD camera (Hamamatsu). Identified cells were visualized with 10x or 40x Zeiss water-immersion objectives. Recordings were performed at 31-34°C.

The base extracellular recording solution contained the following (in mM): 125 NaCl, 25 NaHCO_3_, 1.25 NaH_2_PO_4_, 25 glucose, 3 KCl, 1 MgCl_2_, 2 CaCl_2_ (pH 7.3 and adjusted to 295 mOsm), and was oxygenated (95% O_2_, 5% CO_2_). The pipette solution for whole-cell recordings of excitatory currents contained: 125 K-gluconate, 2 MgCl_2_, 0.025 CaCl_2_, 1 EGTA, 2 Na_3_ATP, 0.5 Na_3_GTP, and 10 HEPES (pH 7.3 with KOH). Recordings of sIPSCs were conducted with equimolar replacement of K-gluconate with cesium methanosulfonate. 100 μM Alexa 488 was added to the pipette solution to allow visualization of cell processes. Patch pipettes, fabricated from borosilicate glass, were pulled to a resistance of 2-3 MΩ for MCs and 4-6 MΩ for eTCs. Current and voltage signals were recorded with a Multiclamp 700B amplifier (Molecular Devices, San Jose, CA), low-pass filtered at 1.8 kHz using an eight-pole Bessel filter, and digitized at 10 kHz. Data were acquired using Axograph X software. Recording sessions with access resistances higher than 15 MΩ were discarded. Reported membrane potential values were corrected for a 7 mV liquid junction potential for our recordings. Epifluorescence was captured by an Axiocam HSm (Zeiss) camera; images were acquired using AxioVision software (Zeiss).

Stimulation of OSN axons in some experiments was performed using a broken-tip patch pipette (5-10 μm-diameter) filled with extracellular solution. The stimulation pipette was placed in the olfactory nerve layer, 50-100 μm superficial to the glomerular layer. Current injections were delivered by a stimulus isolator (World Precision Instruments) under control of a TTL output from Axograph X software. The stimulus interval was 15 seconds.

Cell identity in the recordings was determined in part by visualizing Alexa 488 fluorescence signals. MCs were easily distinguished by their large size and location in the mitral cell layer. eTCs were distinguished from other cells in the glomerular layer by their position in the inner half of the layer, their relatively large, spindle-shaped somas (≥10 μm diameter), a single highly branched apical dendrite that filled most of a glomerulus and no apparent lateral dendrite, and a relatively low input resistance (∼0.2 GΩ; (Hayar et al., 2004)).

### Analysis of electrophysiological data

To evaluate the current variance in the recordings from MCs and eTCs, current traces were analyzed in windows of >3 seconds. To remove the contribution of potential drift in the current to the variance estimate, a sloping baseline was first estimated and subtracted from each trace.

Estimates of MC resting potentials were made immediately after whole-cell break-in. For *Fmr1* KO mice, where MCs engaged in spontaneous LLDs, baseline resting potentials were estimated from periods between the depolarizations. In the analysis of MC spike activity in WT and *Fmr1* KO mice, effort was made to keep the MC baseline potentials near the original estimated resting potentials. In some cases, this required injection of hyperpolarizing current into the cells.

An event detection routine in Axograph was used to detect fast spontaneous excitatory and inhibitory post-synaptic currents (sEPSCs and sIPSCs) in MCs. In the analysis of sEPSCs in MCs, there were in fact few if any events that could be clearly discerned by eye. The reported very low frequencies of sEPSCs reflected a few events that were selected by the event detection algorithm, but these events generally had low amplitude (<20 pA). In the analysis of sIPSCs in MCs, we used a template for the event detection that was similar for WT and *Fmr1* KO mice (decay time constant = 3.5 ms). There have been reports that *Fmr1* KO can induce modest ∼30% changes in the decay kinetics of sIPSCs (Olmos-Serrano et al., 2010), but we verified by eye that the analysis was capturing the large majority of the events for both mouse types. We also found that making modest adjustments in the template kinetics (adjusting the decay time constant by 30%) did not substantially alter the number of detected events.

To evaluate the relationship between spontaneous and evoked excitatory currents in MCs, an estimate of the pre-stimulus current was obtained over a 20-ms window just prior to stimulation of OSNs. An estimate of the evoked current was obtained from the difference between the pre-stimulus current and the current measured 50 ms after stimulation.

### Behavioral procedure and training

Odors were delivered using a computer-controlled olfactometer (Bodyak and Slotnick, 1999) that delivered a known concentration of an odor. Odors were made weekly with high purity (Sigma-Aldrich; vehicle = mineral oil (MO)) to a final volume of 10 ml. Volatilized odors (1/40 dilution with air) were presented to mice through an odor port. We tested the animals’ ability to discriminate between different monomolecular odors as well as between a single component odor and 1:1 mixture that consisted of two odors (see **Table 1**). The total concentration of odors for each experiment was 1% in MO. When mixtures were used, the concentrations of the individual components were reduced to bring the total concentration to 1%.

**Table 1.** Stimulus pairs presented during go/no-go olfactory discrimination task. Each pair of rewarded (S+) and unrewarded (S–) stimuli were presented in the indicated sequence for each *Fmr1* KO and WT mouse used in the study. The first two pairs were presented during the training phase of the study, the last four during the testing phase.

| S+ stimulus | S– stimulus |
| --- | --- |
| Isoamyl acetate (1%) | Mineral oil (MO) |
| 2-heptanone (1%) | MO |
| 2-heptanone (1%) | 3-heptanone (1% in MO) |
| 2-heptanone (1%) | 2-heptanone (0.5%) + 3-heptanone (0.5%) |
| Ethyl acetate (1%) | Propyl acetate (1%) |
| Ethyl acetate (1%) | Ethyl acetate (0.5%) + Propyl acetate (0.5%) |

For preparing mice for the go/no go behavioral task, mice were water-deprived for 1-2 days until they reached 80% of the normal body weight. Behavior training was performed in the olfactometer chamber where mice move freely (Losacco et al., 2020). All mice were first trained to lick the waterspout to obtain water in the presence of odor (1% isoamyl acetate in mineral oil, v/v) in the ‘begin’ task. Subsequently they learned to discriminate 1% isoamyl acetate (S+) versus mineral oil (S-) in the go/no-go discrimination task, followed by learning to discriminate other odor pairs. Data were analyzed for the go/no-go task, but not for the begin task.

In the go/no-go task, mice self-initiated trials by poking their head into the odor delivery port, breaking a photodiode beam. Mice started the trial spontaneously by poking their nose into the odor spout and licking on the lick port. The delivery of the odors was started at a random time starting 1–1.5 s after nose poke and persisted for 2.5 seconds. The mice had to decide to either lick on the waterspout at least once during each of four 0.5-second bins in the 2 second response window to obtain a 10 μl water reward for the S+ odor or refrain from licking for the S– odor. Licks were detected as electrical connectivity between the waterspout and the ground plate on which they stand. The mice learned to refrain from licking for the unrewarded odor due to the unrewarded effort of sustained licking. For correct rejections mice left the spout shortly after the last lick that took place 0.5–1.8 s after odor onset. Performance was evaluated in blocks of 20 trials, with 10 S+ and 10 S– trials presented at random. A mouse was considered proficient at discrimination when they achieved ≥80% correct response rate during at least three blocks per day for two consecutive days.

### Experimental Design and Statistical Analyses

Data were analyzed using Axograph or Prism software (Graph Pad, San Diego, CA). Values are generally expressed as mean ± standard error (SE). Significance was most commonly determined using two-tailed non-parametric tests, either the Wilcoxon signed-rank test or the Mann-Whitney U test. In instances in which more than two comparisons were made at once, the Kruskal-Wallis test was first performed to determine if there were significant differences present within the large dataset. A value of *p* < 0.05 was considered significant (single asterisks in the figures), except if multiple comparisons were made, in which case the Bonferroni correction was applied.

*N*-values reported for electrophysiological experiments refer to the number of test cells, and all statistical comparisons were made based on measurements across different test cells. We also report the number of mice used for slice studies. *N*-values for the behavioral studies reflect the number of test mice.

## Results

To analyze circuit-level effects of *Fmr1* KO in the olfactory bulb, we conducted whole-cell patch-clamp recordings in bulb slices prepared from both KO (hemizygous *Fmr1^–/y^* males) and wild-type (WT) mice. Neural circuit effects of *Fmr1* KO in some other brain systems can vary in younger mice prior to age P21 (Vislay et al., 2013). In our slice studies, we used older mice (P21-P42) to enable comparisons with our behavioral studies that were conducted in older mice (≥P42).

### Fmr1 KO induces spontaneous long-lasting excitatory events in mitral cells

In order to assess whether *Fmr1* KO alters the behavior of neurons in the olfactory bulb, we began with whole-cell patch-clamp recordings from output mitral cells (MCs). In bulb slices from WT mice, MCs display a stereotyped current response to afferent stimulation of olfactory sensory neurons (OSNs) that includes a fast EPSC reflecting direct OSN inputs (the “OSN-EPSC”), as well as a much slower, glomerulus network-driven response that underlies what has been called a long-lasting depolarization (LLD; Carlson et al., 2000; Schoppa and Westbrook, 2001; De Saint Jan et al., 2009; Najac et al., 2011; Gire et al., 2012; Vaaga and Westbrook, 2016). In bulb slices obtained from *Fmr1* KO mice, the basic form of the MC current response to OSN stimulation (at *V*_hold_ = –77 mV) was similar, with components reflecting both the OSN-EPSC and the LLD (see example in **Fig. 1a_i_**). However, a drastic difference was observed under baseline conditions prior to or in the absence of stimulation. MCs from KO mice displayed robust spontaneous, long-lasting inward current events (two example recordings in **Fig. 1a_i_** and **1a**_ii_), which were generally not apparent in WT mice (**Fig. 1b**). The properties of the spontaneous inward current events in MCs in KO mice were not uniform, differing in their frequency of occurrence and time course; in a subset of MCs (example in **Fig. 1a_ii_**), the inward current events occurred often enough that they appeared to be part of regular oscillations. In order to quantify the difference in spontaneous activity between WT and *Fmr1* KO mice, we computed the variance of the entire current trace recorded under baseline conditions, finding that it was on average ∼4-fold larger in *Fmr1* KO versus WT mice (**Fig. 1c**; *Fmr1* KO mean ± SE = 248 ± 39 pA^2^, *n* = 26 cells from 13 mice; WT = 61 ± 12 pA^2^, *n* = 26 cells from 16 mice; *p* < 0.0001 in Mann Whitney *U* test). In spite of these changes in spontaneous prolonged excitatory currents in MCs, *Fmr1* KO did not alter the frequency of kinetically fast sEPSCs that could have reflected direct OSN inputs (*Fmr1* KO: 0.029 ± 0.010 Hz, *n* = 14 cells randomly selected from 26 cell-set; WT: 0.031 ± 0.018 Hz, *n* = 10 cells randomly selected from 26 cell-set; *p* = 0.49 in Mann Whitney test; **Fig. 1d**). In MCs from both WT and *Fmr1* KO mice, these events were very rarely observed, with estimated frequencies that were generally below 0.05 Hz.

**Figure 1.**
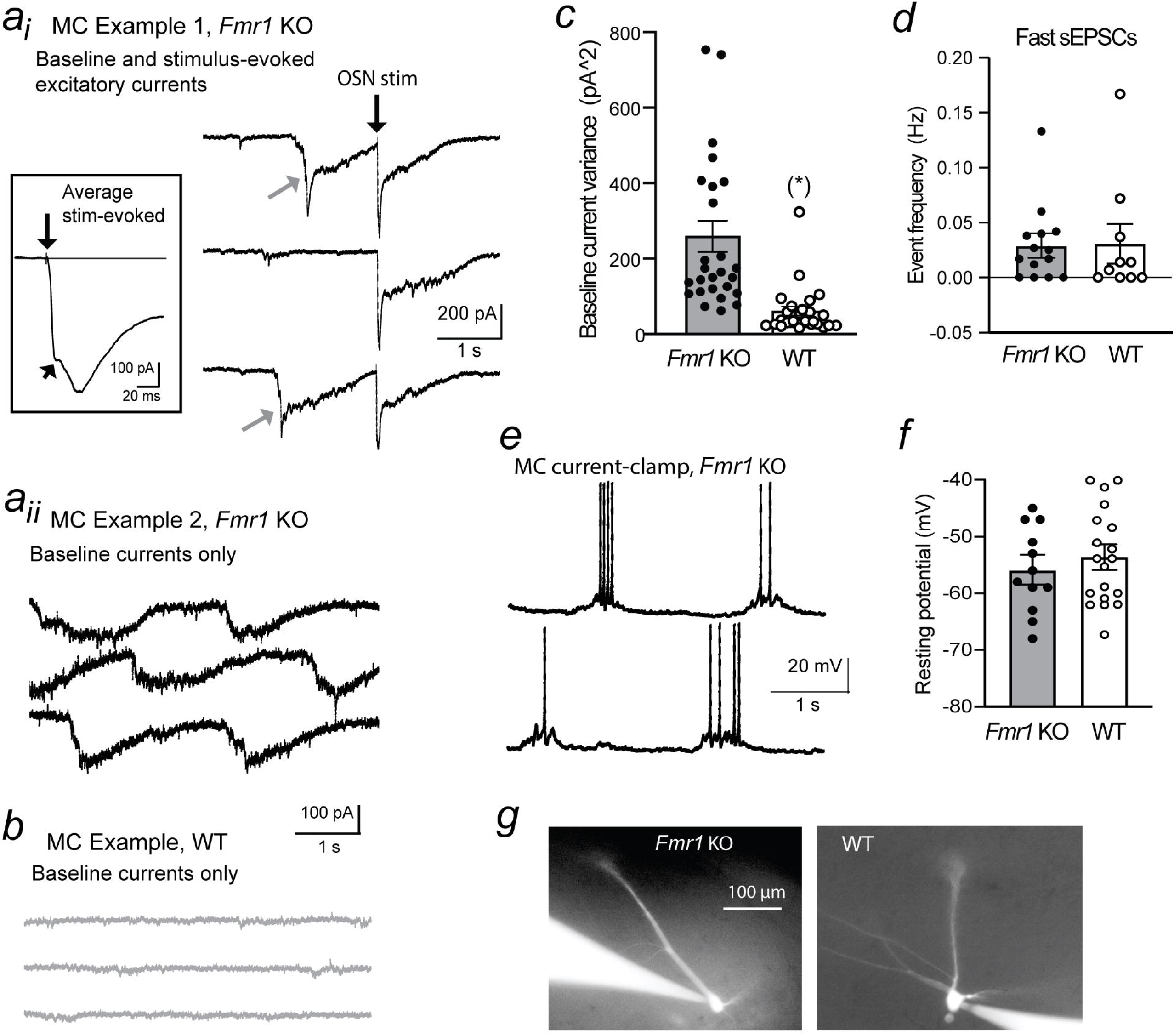
*Fmr1* KO induces spontaneous excitation of MCs. (***a***) Example voltage-clamp recordings (*V*_hold_ = –77 mV) from two mitral cells (MCs; in ***a_i_*** and ***a_ii_***) in bulb slices taken from *Fmr1* KO mice. For the MC in ***a_i_***, electrical stimulation of OSNs (65 µA) was applied. This elicited an excitatory current response with multiple components that included a presumed EPSC reflecting direct OSN inputs (deflection at short diagonal arrow on average trace in inset) along with a complex longer-lasting current response that underlies a long-lasting depolarization (LLD; Carlson et al., 2000). Both MCs displayed numerous spontaneous long-lasting current events that either preceded stimulation (diagonal arrows on raw traces in ***a_i_***) or occurred in the absence of stimulation (in ***a_ii_***). Note that the stimulus-evoked current response in the MC in ***a_i_*** had three components, the OSN-EPSC and fast and slow parts of the LLD. (***b***) Spontaneous long-lasting inward current events were absent in an example recording from a MC in a WT mouse. (***c***) The increase in spontaneous currents in *Fmr1* KO mice was reflected as a large increase in the variance of the baseline current measured in the absence of stimulation (\**p* < 0.0001 in Mann-Whitney *U* test). (***d***) *Fmr1* KO mice did not display an increase in spontaneous fast EPSCs that could have reflected direct OSN inputs. (***e***) In a current-clamp recording from a MC in KO mice, spontaneous phasic depolarizations were observed that induced bursts of action potentials. (***f***) *Fmr1* KO did not alter MC resting potentials. The baseline resting potential for MCs from *Fmr1* KO mice was estimated from periods between the spontaneous long-lasting depolarizations. (***g***) Dye-fills (100 µM Alex 488) of example MCs from *Fmr1* KO and WT mice.

In whole-cell current-clamp recordings from MCs, we found that the spontaneous prolonged inward currents in *Fmr1* KO mice could induce prolonged depolarizations (5.7 ± 0.5 mV, *n* = 5 cells in 3 mice) that had associated bursts of action potentials (**Fig. 1e**). Such spontaneous depolarizations and action potential bursts were not apparent in MCs from WT mice in the absence of stimulation (*n* = 7 cells in 4 mice). The additional spontaneous action potentials in MCs from KO mice occurred in spite of the fact that the resting potentials of MCs in KO mice measured between the prolonged depolarizations (–55.9 ± 2.3 mV, *n* = 12 cells in 5 mice) were not significantly different from the resting potentials in MCs in WT mice (–53.5 ± 1.9 mV, *n* = 19 cells in 12 mice, *p* = 0.57 in Mann-Whitney U test; **Fig. 1f**). This argued that the additional MC action potentials in KO mice could be attributed to the spontaneous depolarizations induced by KO and not due to an overall depolarizing shift in resting potential.

In this initial analysis of MCs in *Fmr1* KO mice, we did not conduct a thorough evaluation of cell anatomy. It was nevertheless notable that the basic MC morphology appeared to be preserved in KO mice (**Fig. 1g**). All of our test MCs in KO mice (*n* = 23) had multiple lateral dendrites, along with one apical dendrite. We did not observe an increase in the number of apical dendrites of MCs. A prior study (Galvez et al., 2005) reported a modest increase in the number of apical dendrites in MCs in *Fmr1* KO mice that were back-crossed into the FVB strain of mice. The same study however reported no effect of KO on dendrite number in the C57BL/6 strain that we used.

### Spontaneous slow excitatory events in MCs from Fmr1 KO mice reflect spontaneous LLDs that originate in glomeruli

We next sought to determine the mechanistic basis for the spontaneous slow excitation in MCs in *Fmr1* KO mice. One clue was provided by a series of recordings from MCs from KO mice that had apical dendrites sectioned prior to reaching the glomerular layer (**Fig. 2a**); this sectioning was fortuitous, accomplished during the preparation of the bulb slices. In none of these MCs (*n* = 11 cells in 5 mice) did we observe slow excitatory current events, a fact that was also reflected in a large reduction in current variance in these MCs (41± 4 pA^2^, *n* = 11) versus those with intact apical dendrites (*p* < 0.0001 in Mann-Whitney U test; **Fig. 2b**). This argued that spontaneous slow excitation originated in the glomerular tuft of MCs. Interestingly, the current variance in MCs with sectioned apical dendrites from KO mice were indistinguishable from variance values in intact MCs from WT mice (*p* = 0.65 in Mann-Whitney *U* test), arguing that the enhanced variance in intact MCs from KO mice (**Fig. 1c**) could be attributed mainly to mechanisms in the MC glomerular tuft. In terms of pharmacological profile, we found that the spontaneous slow excitatory currents in MCs from KO mice were enhanced by the GABA_A_ receptor blocker gabazine (Gbz, 10 µM; 677 ± 286% increase in MC current variance; *p* = 0.0041 in Wilcoxon signed-rank test, *n* = 7 cells in 6 mice; variance in Gbz = 1926 ± 572 pA^2^; **Fig. 2c**) and eliminated by application of ionotropic glutamate receptor blockers 2,3-dioxo-6-nitro-7-sulfamoyl-benzo[f]quinoxaline (NBQX; 10 µM) and DL-2-Amino-5-phosphonopentanoic acid (DL-AP5, 50 µM; MC current variance in drugs = 22 ± 2 pA^2^, *n* = 5 cells in 3 mice). This argued that the excitatory current events in *Fmr1* KO mice were normally down-regulated by GABAergic inhibitory inputs and required glutamate receptor activation.

**Figure 2.**
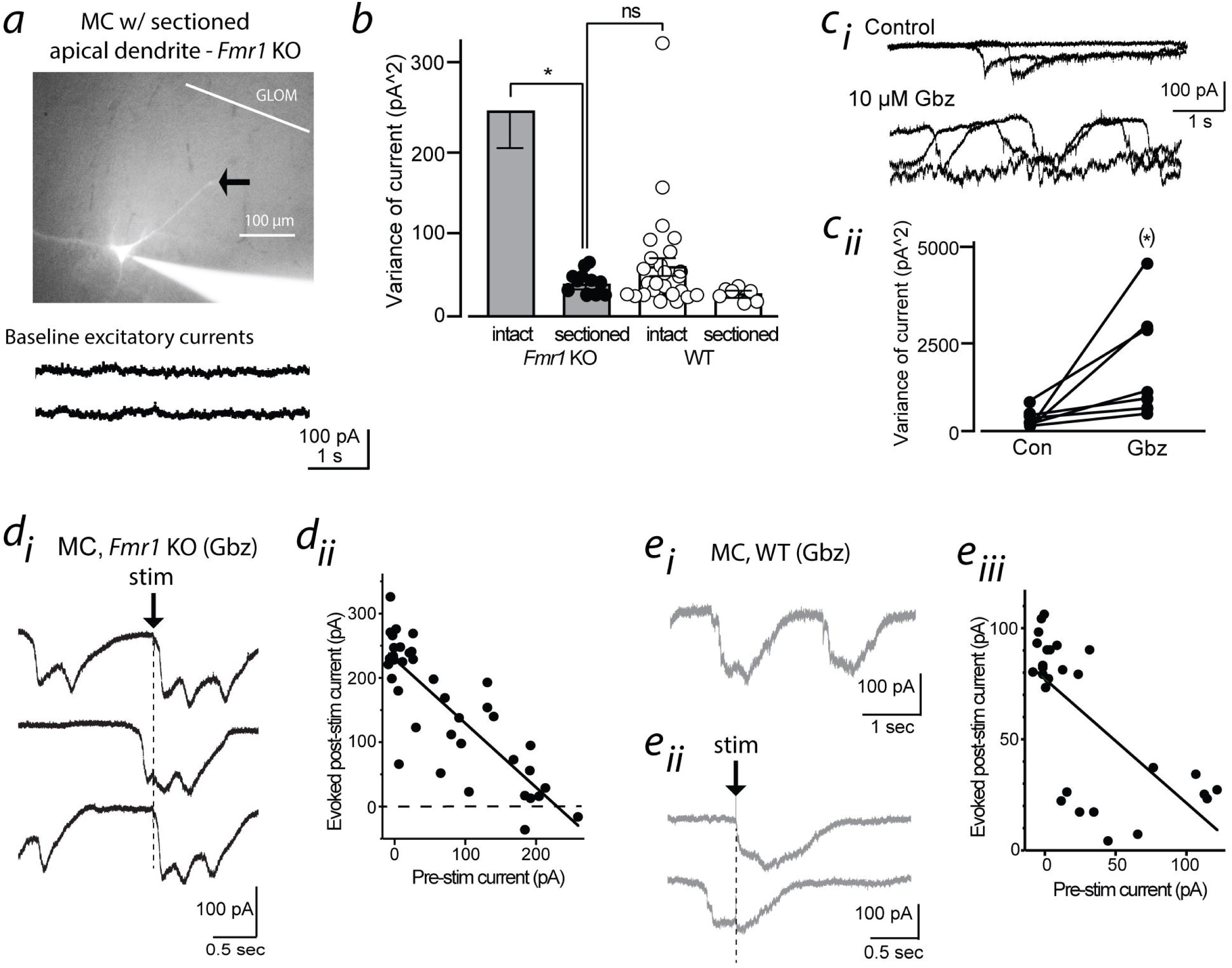
Spontaneous events in MCs from *Fmr1* KO mice reflect currents that underlie long-lasting depolarizations (LLDs) that originate in apical dendrites. (*a*) Example current recording (*V_hold_* = –77 mV) from a MC with sectioned apical dendrite (at arrow in associated image) that lacked spontaneous excitatory currents. (*b*) Summary: dendritic sectioning drastically reduced MC baseline current variance in *Fmr1* KO mice (\**p* < 0.0001 in Mann-Whitney *U* test), bringing it to the level of WT MCs with intact apical dendrites (ns = no significant difference). Plot also shows variance values for MCs from WT mice with sectioned apical dendrites. (*c*) Spontaneous currents in *Fmr1* KO mice were enhanced by block of GABA*_A_* receptors with gabazine (Gbz, 10 µM). Three superimposed raw traces from one MC are shown under each condition (*c_i_*), along with summary results from seven MCs (*c_ii_*; \**p* = 0.0156 in Wilcoxon signed rank test). (*d*) Spontaneous long-lasting events in *Fmr1* KO mice occluded LLDs induced by stimulation of OSNs (100 µA). Occlusion is represented in the middle raw trace in *d_i_* when stimulation elicited no additional current on top of the already-occurring spontaneous event. Occlusion was also reflected in *d_ii_* as a negative correlation (*R^2^* = 0.83, *p* < 0.0001) between the magnitude of pre-stimulus current versus the current evoked 100 ms after the stimulus. Gabazine was added to the bath in these experiments, since the drug significantly increased the probability of sLLDs, making it easier to analyze occlusion across many trials. (*e*) Spontaneous events also occur in WT mice when recordings are conducted in gabazine (10 µM). Example baseline current trace in *e_i_*. Parts *e_ii_* and *e_iii_* show occlusion of stimulus-evoked LLDs (*R^2^* = 0.68, *p* < 0.0001 in *e_iii_*).

We wondered whether the spontaneous slow excitatory events were mechanistically similar to the previously characterized LLDs in MCs that are evoked by OSN stimulation (Carlson et al., 2000; Schoppa and Westbrook, 2001). Both the glomerular origin of the prolonged excitatory events along with their modulation by GABA_A_ receptors fit with prior reports of LLDs. There was also a striking visual resemblance between individual slow excitatory events and stimulus-evoked LLDs in the same MCs (see examples in **Fig. 1a_i_**). That the slow events are the same as stimulus-evoked LLDs was also supported by the fact that spontaneous current events occluded the generation of stimulus-evoked LLDs (**Fig. 2d**). In experiments in which we monitored the timing of both spontaneous and stimulus-evoked events (in the presence of Gbz to increase the frequency of the slow events), we found that stimulation was ineffective at evoking substantial additional current in stimulus trials in which a spontaneous inward current event was already occurring at the point of stimulation (see middle example trace in **Fig. 2d_i_**). Occlusion was also reflected by a strong negative correlation between the magnitude of the pre-stimulus current and the additional current evoked by OSN stimulation (**Fig. 2d_ii_**; *R^2^* = 0.86 ± 0.03, *n* = 4 cells in 2 mice; *p* ≤ 0.00037). Because of these various results, we refer to the spontaneous excitatory events in MCs as spontaneous LLDs (sLLDs) throughout the rest of this study.

As a final point of characterization of the sLLDs, we asked whether the sLLDs could be observed in WT mice under certain conditions. Prior analysis in the rat olfactory bulb (Carlson et al., 2000; Schoppa and Westbrook, 2001) has shown that current events that resemble LLDs can occur spontaneously in wild-type animals when GABA_A_ receptors are blocked. Indeed, we also found this to be the case in our WT mice. When recordings were conducted in the presence of Gbz, MCs in WT mice displayed robust sLLDs (**Fig. 2e_i_**; variance = 941 ± 180 pA^2^, *n* = 7 cells in 7 mice), which, moreover, occluded the generation of stimulus-evoked LLDs (**Fig. 2e_ii_**). The sLLDs in Gbz in WT mice were not significantly different from the sLLDs in KO mice in Gbz (*p* = 0.62 in Mann Whitney *U*-test comparing MC current variance under two conditions). The presence of sLLDs in WT mice in Gbz was important, as they indicated that the basic phenomenon of the sLLDs in MCs is not unique to *Fmr1* KO mice; they also regularly occur in WT mice with relatively moderate changes in recording conditions. In addition, the fact that addition of the GABA_A_ receptor blocker could make MC currents in WT mice resemble currents in KO mice provided one plausible hypothesis for specific network alterations in KO mice, i.e., that KO induced sLLDs by reducing GABAergic inhibition in the bulb.

### Enhanced sLLDs in MCs reflect alterations in the E/I balance at the level of eTCs rather than reduced inhibition on MCs

In order to assess whether GABAergic inhibition in MCs was altered in *Fmr1* KO mice, we next recorded spontaneous inhibitory synaptic activity in MCs. For these experiments, we used a depolarizing holding potential (–7 mV) to help isolate chloride-mediated synaptic currents. We also conducted experiments in the presence of glutamate receptor blockers (10 µM NBQX and 50 µM DL-AP5) to remove any indirect effect on the sIPSCs that resulted secondarily from enhanced spontaneous excitation in KO mice. Under these conditions, we were able to isolate synaptic events with the relatively fast kinetics that are typical for GABA_A_ receptor-mediated events (**Fig. 3a**); these events were also blocked by a GABA_A_ receptor blocker (*n* = 5). In comparisons of the sIPSCs between KO and WT mice, we in fact found no substantial differences. MCs from WT and KO mice had similar sIPSC frequencies (**Fig 3b**; *Fmr1* KO: 1.70 ± 0.40 Hz, *n* = 15 cells in 7 mice; WT: 2.38 ± 0.36 Hz, *n* = 21 cells in 8 mice; *p* = 0.205 in Mann-Whitney *U* test) and amplitude (*Fmr1* KO: 44 ± 3 pA, *n* = 15; WT: 43 ± 3 pA, *n* = 21; *p* = 0.534 in Mann-Whitney *U* test). Moreover, while *Fmr1* KO has been reported to cause modest changes in sIPSC kinetics in other systems (Vislay et al., 2013), the decay kinetics of the sIPSCs were similar for MCs from *Fmr1* KO and WT mice (data not shown; *Fmr1* KO decay time constant: 4.6 ± 0.5 ms, *n* = 15; WT: 4.9 ± 0.3 ms, *n* = 21; *p* = 0.122 in Mann-Whitney *U* test). Thus, contrary to our predictions based on the GABA_A_ receptor pharmacological profile of the sLLDs, *Fmr1* KO did not appear to significantly alter GABAergic synaptic connections onto MCs.

**Figure 3.**
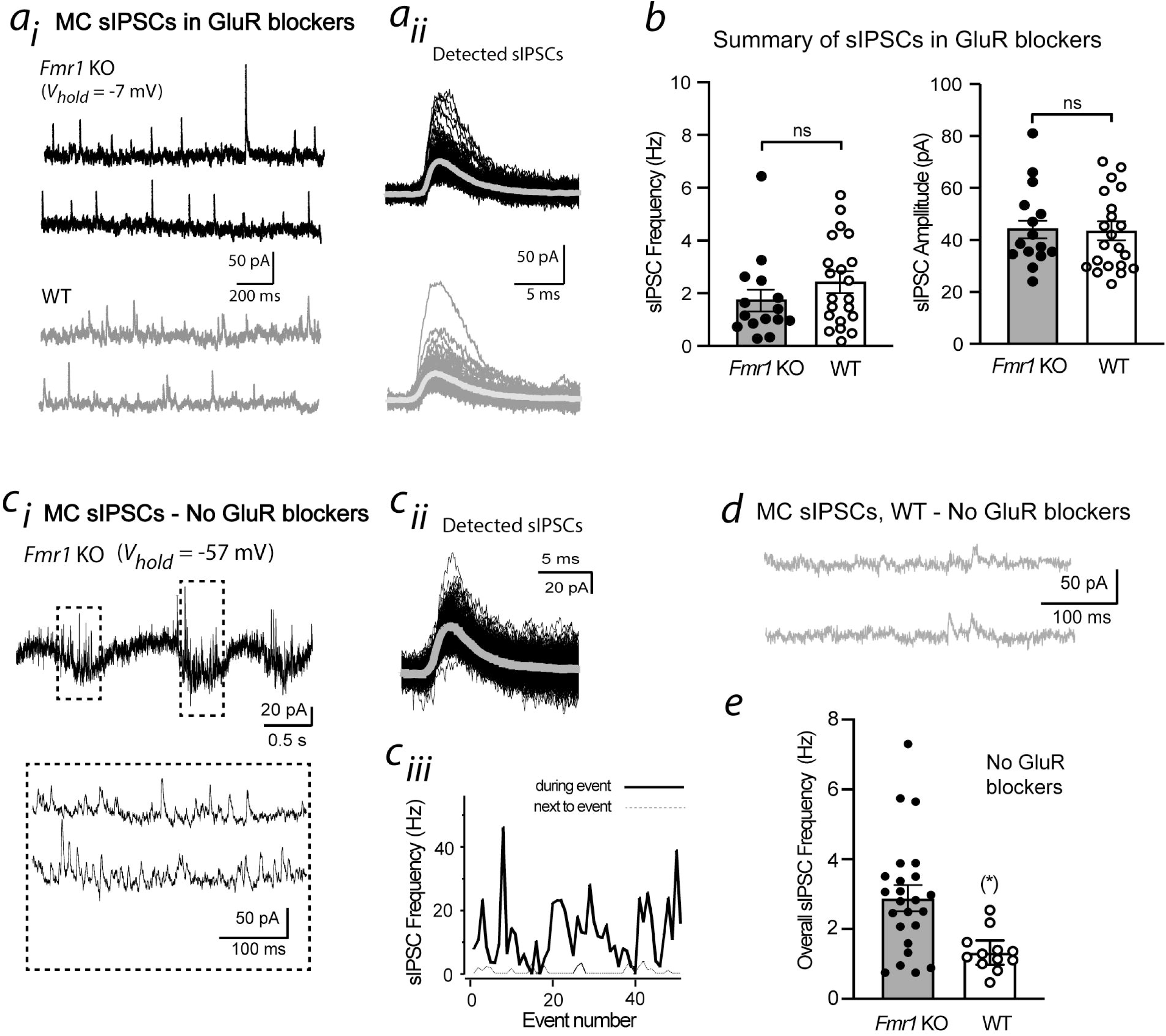
Effects of *Fmr1* KO on spontaneous inhibitory post-synaptic currents (sIPSCs) in MCs. (***a***) Example recordings of sIPSCs in MCs (at *V_hold_* = –7 mV) from *Fmr1* KO (top) and WT (bottom) mice, conducted in the presence of glutamate receptor (GluR) blockers (10 µM NBQX and 50 µM DL-AP5). Raw traces (***a_i_***) and detected sIPSCs (***a_ii_***) are shown for each MC. The thick traces in ***a_ii_*** reflect averages of detected sIPSCs. (***b***) Summary of sIPSC recordings in GluR blockers: *Fmr1* KO did not alter the sIPSC frequency or amplitude. (***c***) Example recording of sIPSCs from a MC in *Fmr1* KO mice made in the absence of GluR blockers (at *V_hold_* = –57 mV). Note in the raw current trace (***c_i_***) the high rate of outward-going sIPSCs associated with each long-lasting inward current event. By analyzing the detected sIPSCs (***c_ii_***), we determined that the sIPSC frequency was much higher during each inward current event versus intervening periods (***c_iii_***). (***d***) Example recording of sIPSCs from a MC in a WT mouse, made in the absence of GluR blockers (at *V_hold_* = – 57 mV). (***e***) Summary: *Fmr1* KO greatly enhanced the sIPSC frequency when recordings were conducted in the absence of GluR blockers (\**p* = 0.020 in Mann-Whitney *U* test). For KO, frequency estimates were obtained from the entire recording, not just during the long-lasting inward current events.

We did also conduct recordings of sIPSCs in MCs in the absence of glutamate receptor blockers. Under these conditions, we found that KO induced a large increase in the frequency of sIPSCs in MCs (**Fig. 3c–e**; *Fmr1* KO = 2.82 ± 0.40 Hz, *n* = 24 cells; WT: 1.29 ± 0.32 Hz, *n* = 12 cells; *p* = 0.020 in Mann-Whitney *U* test). For these recordings, we isolated sIPSCs at –57 mV, which enabled us to record both the inward-going current underlying the sLLDs as well as outward sIPSCs at the same time. This showed that the sIPSC increase was clearly associated with sLLD events (**Fig. 3c_i_**, **Fig. 3c_iii_**), consistent with the enhanced sIPSCs being a secondary result of enhanced excitation of MCs in *Fmr1* KO mice. While the sIPSC recordings in the absence of glutamate receptor blockers could not be used to assess inhibitory synaptic connections onto MCs, the enhanced sIPSCs associated with sLLDs does provide one plausible mechanism for how changes in the bulb circuitry could alter olfactory discrimination capabilities (see *Discussion*). It should be noted that the sIPSC frequency values measured in the absence of GluR blockers (**Fig. 3e**) were not significantly higher than those measured in the presence of the blockers (**Fig. 3b**), as might have been expected. This reflected the fact that the sIPSC recordings in the absence of GluR blockers were conducted at less depolarized holding potentials (resulting in less driving force for inward chloride movement).

The sIPSC recordings in the presence of GluR blockers (**Fig. 3a,b**) provided evidence that the enhanced sLLD probability in KO mice was not due to reduced inhibitory connections onto MCs. As a final test of bulbar mechanisms, we considered an alternate possibility that the enhanced sLLDs could reflect alterations in the balance between excitation and inhibition at the level of a different cell type, the external tufted cells (eTCs). eTCs are well-known to provide feedforward excitation onto MCs (Carlson et al., 2000; Schoppa and Westbrook, 2001; De Saint Jan et al., 2009; Najac et al., 2011; Gire et al., 2012; Vaaga and Westbrook, 2016); also, direct excitation of an eTC can specifically drive MC LLDs. Hence, the enhanced sLLD probability in MCs could reflect enhanced excitation of eTCs. To investigate this hypothesis, we recorded from eTCs at a hyperpolarized voltage (*V*_hold_ = –77 mV) in *Fmr1* KO mice and found that, indeed, large, long-lasting inward currents events could be isolated under baseline conditions that were not apparent in recordings from eTCs in WT mice (**Fig. 4a**). The occurrence of the inward current events in eTCs in KO mice, which resembled the sLLDs in MCs, was associated with a ∼10-fold increase in current variance (*Fmr1* KO = 210 ± 68 pA^2^, *n* = 12 cells in 5 mice; WT: 21 ± 6 pA^2^, *n* = 9 cells in 4 mice; *p* < 0.0001 in Mann Whitney *U* test; **Fig. 4b**). The inward currents in eTCs from KO mice were eliminated by application of glutamate receptor blockers (NBQX, 10 µM, plus DL-AP5, 50 µM; variance = 17 ± 5 pA^2^, *n* = 5 cells in 2 mice; *p* = 0.0013 in Mann Whitney *U* test in comparison with *Fmr1* KO control; **Fig. 4b**), such that the variance of the eTC currents in *Fmr1* KO mice resembled that of WT mice (*p* = 0.454 in Mann Whitney *U* test in comparison to WT). These results suggest that at least one of the circuit-level defects in the bulb that lead to enhanced sLLDs in MCs is enhanced glutamatergic excitation of eTCs that provide feedforward excitation onto MCs (see cartoon in **Fig. 4c**).

**Figure 4.**
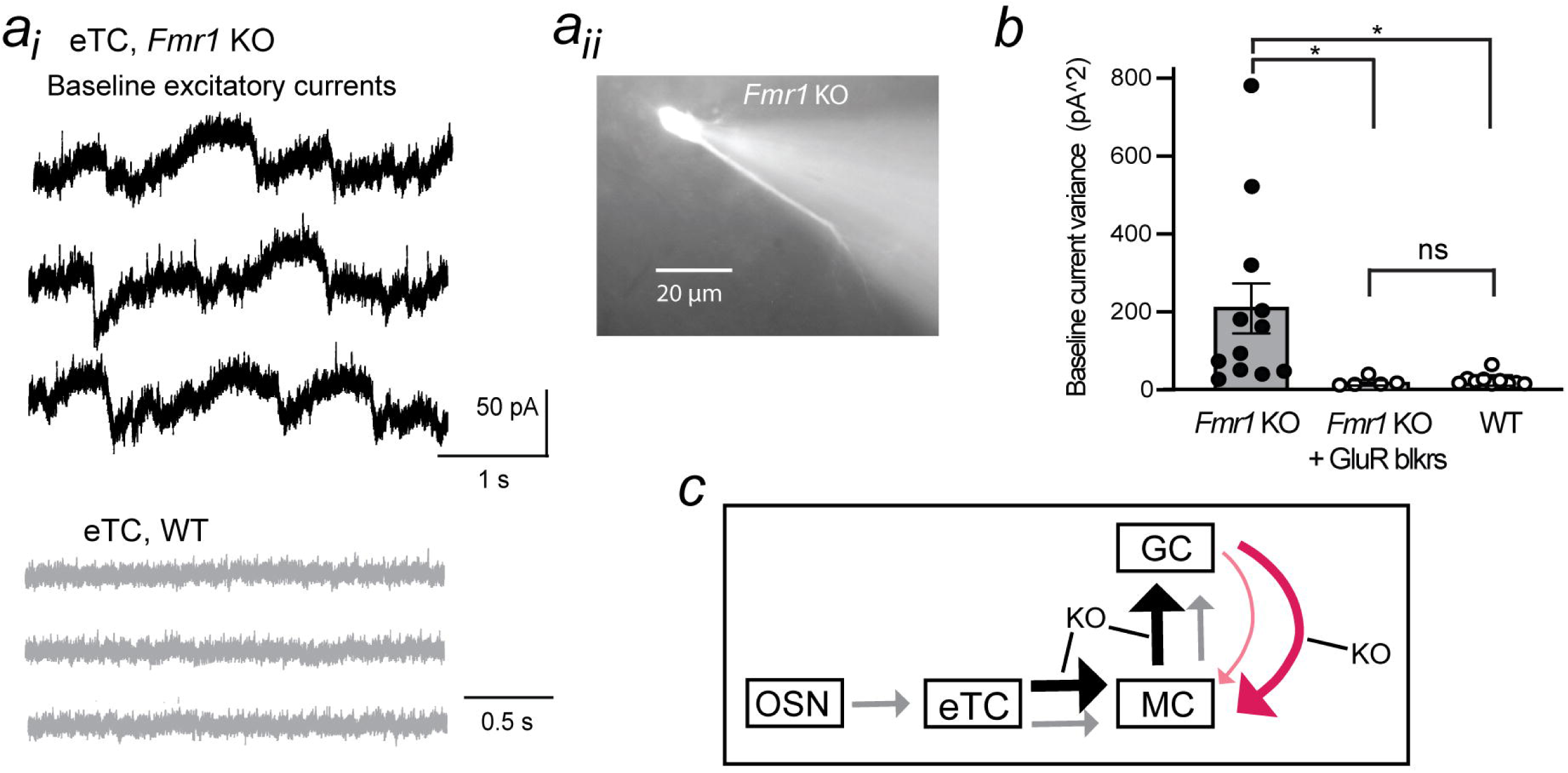
*Fmr1* KO induces spontaneous excitation of eTCs. (***a***) Example voltage-clamp recordings (*V_hold_* = –77 mV) from an eTC in an *Fmr1* KO (top in ***a_i_***) and WT (bottom in ***a_i_***) mouse along with associated fluorescence image of cell (***a_ii_***). Note the repeated occurrence of spontaneous long-lasting inward current events in KO. (***b***) The increase in spontaneous current events in eTCs from *Fmr1* KO mice was reflected in a large increase in the variance of the baseline current (\**p* < 0.0001 in Mann-Whitney *U* test). The histogram also shows that application of GluR blockers (NBQX, 10 µM, plus DL-AP5, 50 µM) greatly reduced variance (\**p* = 0.0013 in Mann-Whitney *U* test), causing eTC currents in KO mice to resemble that of WT mice. (***c***) Model for *Fmr1* KO-induced alterations in the bulb circuit: KO enhances spontaneous excitation of eTCs, which in turn causes enhanced feedforward excitation onto MCs (thick black arrow from eTC to MC). This results in both greater action potential firing in MCs, as well as a secondary increase in transient inhibition (thick red arrow) that is derived from GABAergic (GCs) due to enhanced excitatory drive from MCs. OSN = olfactory sensory neuron.

## *Fmr1* KO mice display deficits in fine odor discrimination

To ascertain the effects of *Fmr1* KO on olfactory behavior, we compared the performance of WT and KO mice in a go/no-go operant discrimination paradigm in which mice were tasked with discriminating two odors, one that was associated with a water reward (S+) and a second (S–) that was not (**Fig. 5a**). In the regimen we used (see sequence of stimuli in **Table 1**), mice first were trained in the go/no-go paradigm wherein they were tasked with discriminating a training odor (1% isoamyl acetate, S+) from mineral oil (MO, S–), followed by discrimination of one of the test odors 2-heptanone (2-Hept, 1% in MO) from MO. Subsequently, mice discriminated various test odor pairs. Responses of a mouse were evaluated in four 20-trial blocks on a given day, with a response counted as correct either when the mouse licked in response to an S+ odor (“Hit”) or when it did not lick in response to an S– odor (“Correct rejection”). Mice were considered proficient in the discrimination of a test odor pair when they achieved ≥80% correct responses in three consecutive blocks on a given day and if they displayed proficiency on two consecutive days. The same seven WT and seven *Fmr1* KO mice were used for all stimulus combinations.

**Figure 5.**
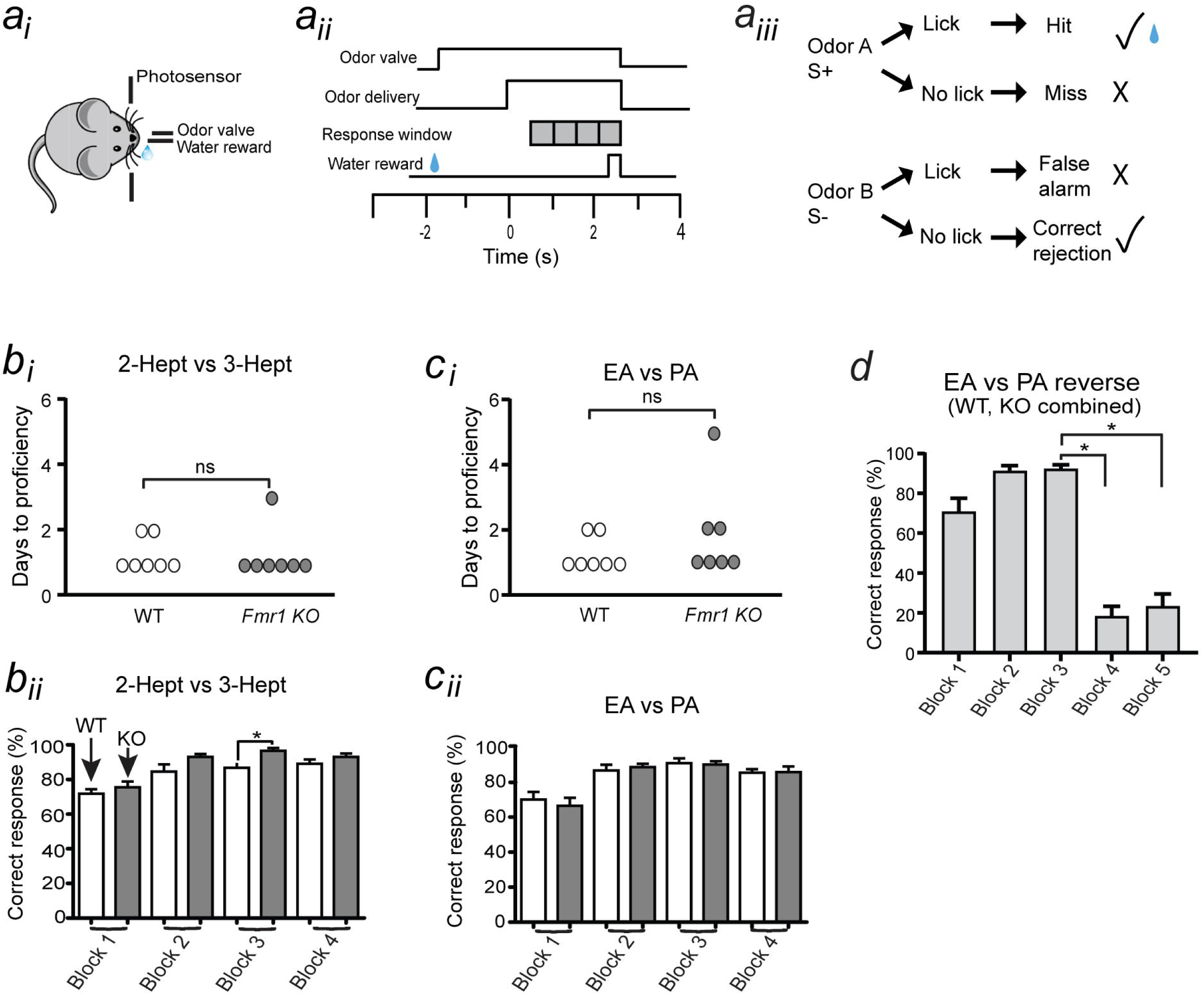
*Fmr1* KO does not alter discrimination of monomolecular odors. (***a***) Go/no-go olfactory discrimination paradigm. (***a_i_***) Mouse self-initiated trials by poking head into the odor delivery port, breaking a photodiode beam. (***a_ii_***) Sequence of steps. When the mouse entered the port, the odor valve opened with odor delivery being initiated 1-1.5 seconds later and lasting for 2.5 seconds. The last 2 seconds of odor delivery was the response window, when the licking of the mouse was assayed to determine whether it was discriminating a rewarded S+ odor from an unrewarded S– odor. The mouse received a water reward if it licked at least once during each of four 0.5-second blocks of the response window upon delivery of the S+ odor. (***a_iii_***) A response was counted as correct (checkmark) if the mouse either licked in response to the S+ odor (Hit) or did not lick in response to S– (Correct rejection). (***b***) *Fmr1* KO and WT mice were similar at discriminating 2-heptanone (2-Hept) from 3-heptanone (3-Hept), as reflected in the fact that the mice took a similar number of days as WT mice to reach proficiency (≥80% correct responses; ***b_i_***) and displayed similar correct response rates upon reaching proficiency (***b_ii_***). Data in ***b_ii_*** reflect correct response rates for each of four consecutive blocks on two days in which proficient performance was displayed. \**p* = 0.0115 in Mann-Whitney *U* test for block 3. (***c***) *Fmr1* KO and WT mice were similar at discriminating ethyl acetate (EA) from propyl acetate (PA). (***d***) For EA versus PA discrimination, an additional day of experiments was added in which the S+ and S– odors were reversed after the third of five blocks. Reversal resulted in a large decrease in correct response rate. \**p* = 0.0020 in Wilcoxon signed-rank test for *Fmr1* KO comparing block 3 with both blocks 4 and 5 (*n* = 6, including 3 WT and 3 KO mice).

In the testing phase, we first compared the ability of WT and *Fmr1* KO mice to discriminate the structurally similar monomolecular odors, 2-heptanone (2-Hept, 1%, S+) and 3-heptanone (3-Hept, 1%, S-). For this pair, we found that WT and KO mice displayed a similar level of discrimination, with each mouse type taking a similar number of days to reach proficiency (**Fig. 5b_i_**; *p* > 0.9999 in Mann Whitney *U* test) and generally displaying similar correct response rates across the four trial blocks upon reaching proficiency (**Fig. 5b_ii_**; see **Table 2** for individual *p*-values). The one exception were values for the correct response rates for block 3, when KO mice performed slightly better than WT (*p* = 0.0115 with Mann-Whitney *U* test). Similar results were obtained when the test odor pairs were ethyl acetate (EA, 1%, S+) and propyl acetate (PA, 1%, S–), both in terms of the number of days to proficiency (**Fig. 5c_i_**; *p* = 0.347 in Mann Whitney *U* test) and correct response rates upon reaching proficiency (**Fig. 5c_ii_**, **Table 2**). For the task involving EA/PA discrimination, an additional day of experiments was added in which the rewarded S+ and unrewarded S– odors were reversed after the third of five blocks (**Fig. 5d**). Reversal resulted in a large decrease in the correct response rate (*p* = 0.0020 in Wilcoxon signed-rank test comparing block 3 and block 4; *n* = 6, combining three WT and three KO mice). This argued that WT and *Fmr1* KO mice were not relying on other cues associated with the S+ and S– stimuli such as auditory cues from the opening of the water reward valve. Taken together, these results imply that *Fmr1* KO does not significantly impact the ability of mice to discriminate some monomolecular odors.

**Table 2.** Statistical comparisons of performance of WT and Fmr1 KO mice in go/no-go discrimination. Mann Whitney *U* and associated *p*-values for various stimulus pairs across four block trials. Analysis was performed on the average values obtained in each block during the last two days that experiments were performed for each stimulus pair. All data reflect comparisons between the same seven WT and seven *Fmr1* KO mice.

| Stimulus pair |  | Block 1 | Block 2 | Block 3 | Block 4 |
| --- | --- | --- | --- | --- | --- |
| Isoamyl acetate 1% vs MO | Mann-Whitney U | 42 | 65 | 57 | 54 |
|  | *p* value | 0.143 | 0.975 | 0.634 | 0.705 |
| 2 Hept 1% vs MO | Mann-Whitney U | 46 | 32 | 45 | 42 |
|  | *p* value | 0.353 | 0.0451 | 0.329 | 0.219 |
| 2 Hept 1% vs 3 Hept 1% | Mann-Whitney U | 42 | 30 | 21 | 41 |
|  | *p* value | 0.372 | 0.0860 | 0.0115 | 0.302 |
| EA 1% vs PA 1% | Mann-Whitney U | 73 | 76 | 79 | 77 |
|  | *p* value | 0.582 | 0.691 | 0.860 | 0.733 |
| 2 Hept (1%) vs 2 Hept (0.5%)<br>– Hept (0.5%) mixture | Mann-Whitney U | 58 | 0 | 0 | 0.500 |
|  | *p* value | 0.636 | <0.0001 | <0.0001 | <0.0001 |
| EA (1%) vs EA (0.5%) – PA<br>(0.5%) mixture | Mann-Whitney U | 24 | 3 | 6.5 | 0 |
|  | *p* value | <0.0001 | <0.0001 | <0.0001 | <0.0001 |

Contrasting results were obtained when the odor discrimination task was made more difficult by having mice discriminate monomolecular odors from odor mixtures. The experiments with mixtures were conducted immediately after the testing phase involving discrimination of monomolecular odors. When mice were tasked with discriminating either 2-Hept (1%, S+) from a 1:1 mixture of 2-Hept and 3-Hept (0.5% for both, S–) or EA (1%, S+) from a 1:1 EA/PA mixture (0.5% for both, S–), WT but not *Fmr1* KO mice were successful in discriminating the stimuli (**Fig. 6a,b**). Correct response rates for *Fmr1* KO mice were much lower than for WT mice for nearly all blocks for both types of mixture experiments (see *p*-values in **Table 2**), remaining near 50% chance level. The poorer performance of KO mice in the mixture experiments occurred in spite of the fact that *Fmr1* KO mice were generally given one additional day of testing versus WT mice (mean days of testing for the two mixture experiments: 2.3 ± 0.2 days for WT, *n* = 14; 3.0 ± 0.0 days for *Fmr1* KO, *n* = 14; *p* < 0.0001 in Mann Whitney *U* test). Importantly, in the mixture experiments involving EA/PA, we found that *Fmr1* KO mice, after just failing at discriminating EA from EA/PA mixture, were successful at discriminating EA (1%, S+) from EA (1%, S–) upon an additional day of retesting (**Fig. 6c**; *p* = 0.0313 in Wilcoxon signed-rank test comparing block 4 of retest day with block 4 of last day of mixture experiment, *n* = 4 KO mice). This argued that the impaired performance of KO mice in discriminating the monomolecular odors from the mixtures was not due to fatigue or loss of interest in the task. We also evaluated whether the incorrect responses for *Fmr1* KO animals in the mixture experiments occurred more often in S+ versus S– trials, finding many more errors for S– trials (**Fig. 6d**; *p* < 0.0001 in Mann Whitney U test for blocks 2, 3, and 4). The higher error rate in the S– trials, when mice licked in response to the unrewarded odor, would be expected due to the water-deprived state of the mice.

**Figure 6.**
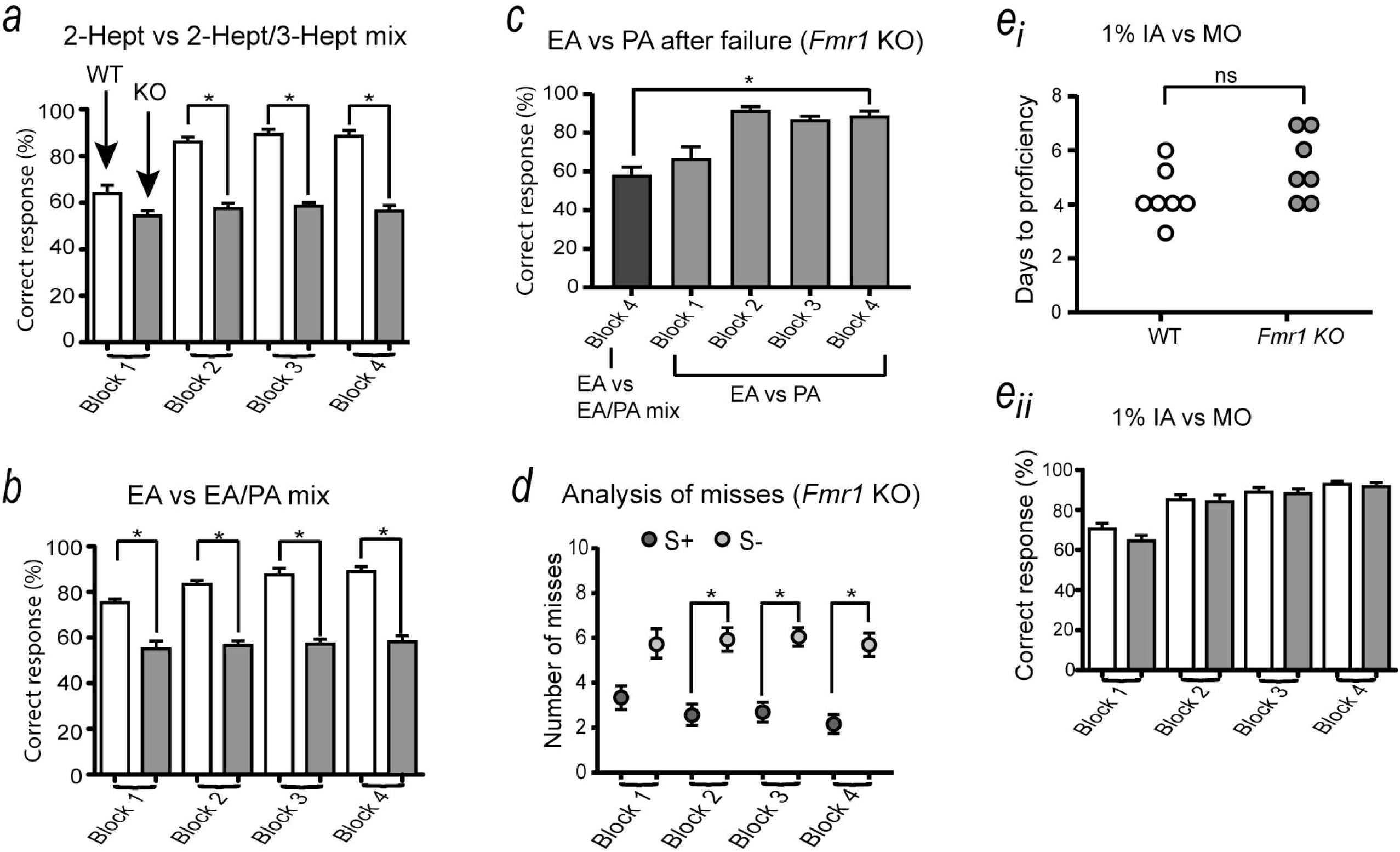
*Fmr1* KO mice display deficits in more difficult discrimination tasks involving odor mixtures. (***a***) Correct response rates when mice were discriminating 2-Hept from a 1:1 mixture of 2-Hept and 3-Hept. Note that, except for the first block, WT mice performed much better than *Fmr1* KO mice. \**p* < 0.0001 in Mann-Whitney *U* test for blocks 2, 3, and 4. Data for both WT and *Fmr1* KO mice reflect the last two days in which experiments were performed with this odor combination; *Fmr1* KO mice were generally given one additional day beyond what WT mice took to reach proficiency (see main text). (***b***) *Fmr1* KO mice performed worse than WT mice in discrimination of EA from a 1:1 mixture of EA and PA. \**p* < 0.0001 in Mann Whitney *U* test for all blocks. (***c***) *Fmr1* KO mice that failed to discriminate EA from the EA/PA mixture were successful at discriminating EA from PA upon retesting. Data reflect four KO mice for block 4 on the last day of testing EA versus EA/PA mixture discrimination (left bar) and four consecutive blocks on a subsequent day of retesting with EA versus PA. \**p* = 0.0313 in Wilcoxon signed-rank test comparing block four of the two days. (***d***) Most mistakes made by *Fmr1* KO mice were during S– trials rather than S+ trials. The plotted data reflect the number of misses made in the half (10) of the 20 trials in each block that were either S– or S+. \**p* < 0.0001 in Mann-Whitney *U* test for blocks 2, 3, and 4. (***e***) Evidence that impaired odor discrimination in *Fmr1* KO mice was not due to cognitive impairments. In the training phase involving discrimination of isoamyl acetate (IA) from mineral oil (MO), KO mice took a similar number of days to reach proficiency (***e_i_***) and displayed similar correct response rates upon reaching proficiency (***e_ii_***). Data in ***e_ii_*** reflect the last two days of experiments with the IA/MO pairs.

One interpretation of the impaired performance of *Fmr1* KO mice in the discrimination experiments involving odor mixtures is that KO mice have olfactory sensory processing deficits, but we also considered whether the observed results could have been explained by general cognitive or motor deficits in KO mice. While these latter mechanisms were difficult to exclude completely, evidence against them being important in our experiments was the fact that *Fmr1* KO mice performed just as well as WT mice in monomolecular odor discrimination (**Fig. 5b,c**). In addition, when we analyzed the performance of KO mice during the initial training phase of the experiments, involving discrimination of isoamyl acetate from MO, we found no differences: *Fmr1* KO and WT mice took a similar number of days to reach proficiency (**Fig. 6e_i_**; *p* = 0.0813 in Mann Whitney *U* test) and displayed similar correct response rates upon reaching proficiency (**Fig. 6e_ii_**; see *p* values in **Table 2**). These results suggest that the reduced performance of *Fmr1* KO mice in the go/no-go discrimination experiments involving odor mixtures was due to sensory processing deficits.

## Discussion

In this study, we examined olfactory circuit and behavioral properties of the *Fmr1* KO mouse model of FXS. Our main findings were that KO induced a large increase in the excitation of bulbar MCs under baseline conditions, i.e., in the absence of a stimulus, and that this was likely at least in part due to enhanced excitation of eTCs. In addition, *Fmr1* KO impaired fine odor discrimination capabilities. Here, we discuss these various results and also how they may relate to each other.

### Deficits in olfactory bulb circuitry due to Fmr1 KO

Evidence for disrupted olfactory bulb circuit properties was obtained first in whole-cell patch-clamp recordings from MCs. MCs from KO mice displayed a massive increase in spontaneous, long-lasting excitatory current events that were largely not present in WT mice. The events, which originated in glomeruli, also were associated with prolonged depolarizations and an increase in spontaneous action potential firing. Several lines of evidence suggest that the excitatory events were spontaneous versions of the well-characterized LLD events that dominate the MC response to stimulation of OSNs (hence, sLLDs; (Carlson et al., 2000; Schoppa and Westbrook, 2001; Vaaga and Westbrook, 2016). These included the slow time course of the spontaneous events, their glomerular origin, their modulation by GABA_A_ receptor antagonists, as well as the fact that the spontaneous currents occluded generation of stimulus-evoked LLDs. One prior study (De Saint Jan et al., 2009) reported spontaneous long-lasting excitatory events in MCs of WT mice, different from our results indicating that such activity was uncommon in control mice. It is unclear what these different results reflect. The important point is that, using identical experimental conditions, we observed a large increase in sLLDs in *Fmr1* KO versus WT mice.

LLDs are network-level events that occur probabilistically depending on the balance between excitation and GABAergic inhibition within a glomerulus (Carlson et al., 2000; Schoppa and Westbrook, 2001; Gire and Schoppa, 2009). Here we were able to identify one likely mechanism whereby *Fmr1* KO increased the probability of sLLDs. Based on recordings on eTCs, it appears that KO caused a massive increase in the spontaneous excitation of eTCs, which are a glutamatergic neuron in the bulb that provides feedforward excitation onto MCs within glomeruli (De Saint Jan et al., 2009; Najac et al., 2011; Gire et al., 2012). Moreover, because excitation of eTCs can directly drive LLDs in MCs (De Saint Jan et al., 2009; Gire et al., 2012), an increase in their spontaneous excitation should naturally increase the probability of sLLDs in MCs. While we did not investigate in detail the nature of the excitatory events in eTCs, they most likely reflected spontaneous versions of LLD events that are known to occur in eTCs as well as MCs (Hayar et al., 2005; Gire and Schoppa, 2009). Our studies also argued against several other candidate mechanisms that could have enhanced the probability of sLLDs in MCs. Most notably, we found, based on recordings of sIPSCs in glutamate receptor blockers, that KO did not alter GABAergic synaptic connections onto MCs. Reductions in inhibitory connections are a commonly observed mechanism by which *Fmr1* KO alters the local excitation/inhibition balance in other brain circuits (Cea-Del Rio and Huntsman, 2014; Takarae and Sweeney, 2017; Telias, 2019).

Studies in mouse models of the broad class of ASDs have begun to identify both physiological and behavioral changes that could be associated with ASDs (Geramita et al., 2020; Li et al., 2023). However, prior analyses of olfactory circuit deficits in mouse models of FXS have been limited. The most extensive such study was conducted by Scotto-Lomassese and co-workers (Scotto-Lomassese et al., 2011), who reported that *Fmr1* KO caused moderate increases in the number and length of dendritic spines on adult-born GABAergic granule cells, although glutamatergic synapses onto the spines were normal. Here, we have for the first time analyzed the physiological properties of output MCs in *Fmr1* KO mice and have found dramatic changes in excitatory events (the LLDs) that reflect the summed contribution of a glomerular network.

Certainly, there are a number of important unresolved mechanistic issues, perhaps the most important of which is exactly how *Fmr1* KO enhances spontaneous excitation of eTCs. One obvious candidate is that KO enhances eTC excitation by reducing inhibitory synaptic connections onto eTCs (coming from periglomerular cells). We found in our pharmacological analysis that block of GABA_A_ receptors can induce sLLDs in MCs that are normally not present in WT mice. This argued that KO could be enhancing the probability of MC sLLDs by reducing GABAergic synapses somewhere in the bulbar circuit. Another possibility is that the enhanced excitation of eTCs in KO mice could reflect increases in persistent sodium currents that play a critical role in regulating eTC excitation (Liu and Shipley, 2008). *Fmr1* KO-induced enhancement of such currents has been reported in pyramidal cells of the entorhinal cortex (Deng and Klyachko, 2016). Future studies in mutant mice in which *Fmr1* is conditionally knocked out in specific cells (Mientjes et al., 2006) will be useful for further interrogating the precise loci of KO-induced alterations in the bulb. It should be noted that enhanced excitation of eTCs is very likely not a result of greater excitation of MCs, since signaling from MCs to eTCs appears to be negligible (De Saint Jan et al., 2009; Gire et al., 2012).

### Deficits in olfactory behavior due to Fmr1 KO

In our analysis of olfactory behavior, we found that *Fmr1* KO mice displayed significant deficits in a go/no-go olfactory discrimination task in which the mice were tasked with distinguishing a monomolecular odor from an odor mixture. In contrast, KO mice performed as well as WT mice in an easier task involving discrimination of two similar monomolecular odors. In these latter experiments, the mice required a similar number of days to reach proficiency after initiating the task and displayed similar levels of proficiency. This was important for two reasons. First, it suggested that *Fmr1* KO mice are generally quite good at discriminating odors, only showing impairments for difficult tasks. In addition, the results with monomolecular odor discrimination serve as a reasonable control for other possible explanations for the impaired performance in fine odor discrimination task for the KO mice other than sensory deficits. For example, if the impaired performance in the mixture experiments were due to cognitive or motor deficits, the impairments likely would have appeared in the easier discrimination task.

Only a few prior studies have examined olfactory behavioral deficits in *Fmr1* KO mice, with somewhat conflicting results. Schilit Nitenson and co-workers (Schilit Nitenson et al., 2015) provided evidence, using an olfactory cross-habituation task, that KO reduces olfactory sensitivity, although they did not find impairments in olfactory discrimination. In contrast, Larson and co-workers (Larson et al., 2008) found impaired olfactory discrimination in KO mice with the go/no-go task, both slower learning and increases in failure to discriminate, but no differences in olfactory sensitivity. Certainly at least some of differences in these results may have been explained by the different behavioral tests, but the results of our behavioral studies here also point to the importance of task difficulty, i.e., the odor pairs used, in what results might be expected in experiments that assess odor discrimination deficits. Task difficulty was not systematically accounted for in the prior studies. As for *Fmr1* KO effects on olfactory sensitivity, we did not examine this issue here, opting instead to perform a more detailed analysis of odor discrimination.

If we consider the observed deficits in fine odor discrimination in *Fmr1* KO mice, how might these be related to the circuit-level effects of KO on spontaneous excitation of MCs that we also observed (**Fig. 1**)? One model builds on the common assumption that animals discriminate odors based on differences in which subpopulations of MCs are activated by those odors (Yokoi et al., 1995). The higher propensity for MCs in KO mice to fire action potentials spontaneously could interfere with detection of those differences, since any spontaneous activity that co-occurs with odor-evoked activity would necessarily not be odor-specific. It is also possible that what is most functionally important is not the enhanced spontaneous activity in MCs per se but rather a generally higher tendency for MCs to be excited. MCs in a more excitable state should naturally be more easily excited by odors, which in turn should result in a broadening of MC odor-tuning curves. Broader neural tuning has already been implicated as being at least correlated with reductions in odor discrimination capabilities in a *Drosophila* model of FXS (Franco et al., 2017) and also may have a role in tactile impairments in *Fmr1* KO mice (Arnett et al., 2014; Juczewski et al., 2016; Franco et al., 2017).

Studies across animal species have also suggested that odor discrimination might depend on the properties of gamma frequency (40-100 Hz in mammals) synchronized oscillations that are evoked by odors (Stopfer et al., 1997; Beshel et al., 2007; Lepousez and Lledo, 2013; Kay, 2015; Losacco et al., 2020). Both eliminating and enhancing gamma oscillations can impair odor discrimination (Stopfer et al., 1997; Lepousez and Lledo, 2013; Kay, 2015), which suggests that there is an optimal level of gamma oscillations for discrimination. In the present study, we did not test directly whether *Fmr1* KO mice have altered gamma oscillations. However, we did observe that KO increased transient inhibitory synaptic activity that underlie gamma oscillations (Bazhenov et al., 2001; Galan et al., 2006; Schoppa, 2006), when sIPSC recordings were performed in the absence of GluR blockers (**Fig. 3c–e**). The enhanced sIPSCs in these experiments resulted from enhanced excitatory drive onto GABAergic neurons in KO mice (**Fig. 4c**). Future *in vivo* studies should shed light onto whether *Fmr1* KO indeed alters gamma oscillations and how this might interfere with odor discrimination.

## Acknowledgements

This work was supported by NIH grants R01DC006640 (NES) and F31DC017350. We thank Dr. Diego Restrepo (UCAMC) and personnel in his lab for advice and assistance in training Dr. Kuruppath in the go/no-go olfactory discrimination behavioral task.

## References

Akins MR, Leblanc HF, Stackpole EE, Chyung E, Fallon JR (2012) Systematic mapping of fragile X granules in the mouse brain reveals a potential role for presynaptic FMRP in sensorimotor functions. The Journal of comparative neurology 520:3687–3706.

Arnett MT, Herman DH, McGee AW (2014) Deficits in tactile learning in a mouse model of fragile X syndrome. PloS one 9:e109116.

Bagni C, Zukin RS (2019) A Synaptic Perspective of Fragile X Syndrome and Autism Spectrum Disorders. Neuron 101:1070–1088.

Bazhenov M, Stopfer M, Rabinovich M, Huerta R, Abarbanel HD, Sejnowski TJ, Laurent G (2001) Model of transient oscillatory synchronization in the locust antennal lobe. Neuron 30:553–567.

Beshel J, Kopell N, Kay LM (2007) Olfactory bulb gamma oscillations are enhanced with task demands. The Journal of neuroscience: the official journal of the Society for Neuroscience 27:8358–8365.

Bhattacharyya U, Bhalla US (2015) Robust and Rapid Air-Borne Odor Tracking without Casting. eNeuro 2.

Bodaleo F, Tapia-Monsalves C, Cea-Del Rio C, Gonzalez-Billault C, Nunez-Parra A (2019) Structural and Functional Abnormalities in the Olfactory System of Fragile X Syndrome Models. Front Mol Neurosci 12:135.

Bodyak N, Slotnick B (1999) Performance of mice in an automated olfactometer: odor detection, discrimination and odor memory. Chemical senses 24:637–645.

Boudjarane MA, Grandgeorge M, Marianowski R, Misery L, Lemonnier E (2017) Perception of odors and tastes in autism spectrum disorders: A systematic review of assessments. Autism Res 10:1045–1057.

Brackett DM, Qing F, Amieux PS, Sellers DL, Horner PJ, Morris DR (2013) FMR1 transcript isoforms: association with polyribosomes; regional and developmental expression in mouse brain. PloS one 8:e58296.

Carlson GC, Shipley MT, Keller A (2000) Long-lasting depolarizations in mitral cells of the rat olfactory bulb. The Journal of neuroscience: the official journal of the Society for Neuroscience 20:2011–2021.

Cea-Del Rio CA, Huntsman MM (2014) The contribution of inhibitory interneurons to circuit dysfunction in Fragile X Syndrome. Frontiers in cellular neuroscience 8:245.

Chen L, Toth M (2001) Fragile X mice develop sensory hyperreactivity to auditory stimuli. Neuroscience 103:1043–1050.

Christie SB, Akins MR, Schwob JE, Fallon JR (2009) The FXG: a presynaptic fragile X granule expressed in a subset of developing brain circuits. The Journal of neuroscience: the official journal of the Society for Neuroscience 29:1514–1524.

Davis JK, Broadie K (2017) Multifarious Functions of the Fragile X Mental Retardation Protein. Trends Genet 33:703–714.

De Saint Jan D, Hirnet D, Westbrook GL, Charpak S (2009) External tufted cells drive the output of olfactory bulb glomeruli. The Journal of neuroscience: the official journal of the Society for Neuroscience 29:2043–2052.

Deng PY, Klyachko VA (2016) Increased Persistent Sodium Current Causes Neuronal Hyperexcitability in the Entorhinal Cortex of Fmr1 Knockout Mice. Cell reports 16:3157–3166.

Dudova I, Vodicka J, Havlovicova M, Sedlacek Z, Urbanek T, Hrdlicka M (2011) Odor detection threshold, but not odor identification, is impaired in children with autism. Eur Child Adolesc Psychiatry 20:333–340.

Franco LM, Okray Z, Linneweber GA, Hassan BA, Yaksi E (2017) Reduced Lateral Inhibition Impairs Olfactory Computations and Behaviors in a Drosophila Model of Fragile X Syndrome. Current biology: CB 27:1111–1123.

Galan RF, Fourcaud-Trocme N, Ermentrout GB, Urban NN (2006) Correlation-induced synchronization of oscillations in olfactory bulb neurons. The Journal of neuroscience: the official journal of the Society for Neuroscience 26:3646–3655.

Galvez R, Smith RL, Greenough WT (2005) Olfactory bulb mitral cell dendritic pruning abnormalities in a mouse model of the Fragile-X mental retardation syndrome: further support for FMRP’s involvement in dendritic development. Brain Res Dev Brain Res 157:214–216.

Geramita MA, Wen JA, Rannals MD, Urban NN (2020) Decreased amplitude and reliability of odor-evoked responses in two mouse models of autism. Journal of neurophysiology 123:1283–1294.

Gire DH, Schoppa NE (2009) Control of on/off glomerular signaling by a local GABAergic microcircuit in the olfactory bulb. The Journal of neuroscience: the official journal of the Society for Neuroscience 29:13454–13464.

Gire DH, Kapoor V, Arrighi-Allisan A, Seminara A, Murthy VN (2016) Mice Develop Efficient Strategies for Foraging and Navigation Using Complex Natural Stimuli. Current biology: CB 26:1261–1273.

Gire DH, Franks KM, Zak JD, Tanaka KF, Whitesell JD, Mulligan AA, Hen R, Schoppa NE (2012) Mitral cells in the olfactory bulb are mainly excited through a multistep signaling path. The Journal of neuroscience: the official journal of the Society for Neuroscience 32:2964–2975.

Hagerman RJ, Berry-Kravis E, Hazlett HC, Bailey DB, Jr., Moine H, Kooy RF, Tassone F, Gantois I, Sonenberg N, Mandel JL, Hagerman PJ (2017) Fragile X syndrome. Nat Rev Dis Primers 3:17065.

Hayar A, Shipley MT, Ennis M (2005) Olfactory bulb external tufted cells are synchronized by multiple intraglomerular mechanisms. The Journal of neuroscience: the official journal of the Society for Neuroscience 25:8197–8208.

Hayar A, Karnup S, Shipley MT, Ennis M (2004) Olfactory bulb glomeruli: external tufted cells intrinsically burst at theta frequency and are entrained by patterned olfactory input. The Journal of neuroscience: the official journal of the Society for Neuroscience 24:1190–1199.

Hinds HL, Ashley CT, Sutcliffe JS, Nelson DL, Warren ST, Housman DE, Schalling M (1993) Tissue specific expression of FMR-1 provides evidence for a functional role in fragile X syndrome. Nat Genet 3:36–43.

Ishii KK, Touhara K (2019) Neural circuits regulating sexual behaviors via the olfactory system in mice. Neurosci Res 140:59–76.

Juczewski K, von Richthofen H, Bagni C, Celikel T, Fisone G, Krieger P (2016) Somatosensory map expansion and altered processing of tactile inputs in a mouse model of fragile X syndrome. Neurobiol Dis 96:201–215.

Kay LM (2015) Olfactory system oscillations across phyla. Current opinion in neurobiology 31:141–147.

Koehler L, Fournel A, Albertowski K, Roessner V, Gerber J, Hummel C, Hummel T, Bensafi M (2018) Impaired Odor Perception in Autism Spectrum Disorder Is Associated with Decreased Activity in Olfactory Cortex. Chemical senses 43:627–634.

Korsak LIT, Shepard KA, Akins MR (2017) Cell type-dependent axonal localization of translational regulators and mRNA in mouse peripheral olfactory neurons. The Journal of comparative neurology 525:2202–2215.

Lai A, Valdez-Sinon AN, Bassell GJ (2020) Regulation of RNA granules by FMRP and implications for neurological diseases. Traffic 21:454–462.

Lane AE, Molloy CA, Bishop SL (2014) Classification of children with autism spectrum disorder by sensory subtype: a case for sensory-based phenotypes. Autism Res 7:322–333.

Larson J, Kim D, Patel RC, Floreani C (2008) Olfactory discrimination learning in mice lacking the fragile X mental retardation protein. Neurobiol Learn Mem 90:90–102.

Lepousez G, Lledo PM (2013) Odor discrimination requires proper olfactory fast oscillations in awake mice. Neuron 80:1010–1024.

Li Y, Swerdloff M, She T, Rahman A, Sharma N, Shah R, Castellano M, Mogel D, Wu J, Ahmed A, San Miguel J, Cohn J, Shah N, Ramos RL, Otazu GH (2023) Robust odor identification in novel olfactory environments in mice. Nature communications 14:673.

Liu S, Shipley MT (2008) Multiple conductances cooperatively regulate spontaneous bursting in mouse olfactory bulb external tufted cells. The Journal of neuroscience: the official journal of the Society for Neuroscience 28:1625–1639.

Losacco J, Ramirez-Gordillo D, Gilmer J, Restrepo D (2020) Learning improves decoding of odor identity with phase-referenced oscillations in the olfactory bulb. eLife 9.

Mientjes EJ, Nieuwenhuizen I, Kirkpatrick L, Zu T, Hoogeveen-Westerveld M, Severijnen L, Rife M, Willemsen R, Nelson DL, Oostra BA (2006) The generation of a conditional Fmr1 knock out mouse model to study Fmrp function in vivo. Neurobiol Dis 21:549–555.

Mila M, Alvarez-Mora MI, Madrigal I, Rodriguez-Revenga L (2018) Fragile X syndrome: An overview and update of the FMR1 gene. Clin Genet 93:197–205.

Najac M, De Saint Jan D, Reguero L, Grandes P, Charpak S (2011) Monosynaptic and polysynaptic feed-forward inputs to mitral cells from olfactory sensory neurons. The Journal of neuroscience: the official journal of the Society for Neuroscience 31:8722–8729.

Olmos-Serrano JL, Paluszkiewicz SM, Martin BS, Kaufmann WE, Corbin JG, Huntsman MM (2010) Defective GABAergic neurotransmission and pharmacological rescue of neuronal hyperexcitability in the amygdala in a mouse model of fragile X syndrome. The Journal of neuroscience: the official journal of the Society for Neuroscience 30:9929–9938.

Rais M, Binder DK, Razak KA, Ethell IM (2018) Sensory Processing Phenotypes in Fragile X Syndrome. ASN Neuro 10:1759091418801092.

Razak KA, Binder DK, Ethell IM (2021) Neural Correlates of Auditory Hypersensitivity in Fragile X Syndrome. Front Psychiatry 12:720752.

Rotschafer SE, Razak KA (2014) Auditory processing in fragile x syndrome. Frontiers in cellular neuroscience 8:19.

Rozenkrantz L, Zachor D, Heller I, Plotkin A, Weissbrod A, Snitz K, Secundo L, Sobel N (2015) A Mechanistic Link between Olfaction and Autism Spectrum Disorder. Current biology: CB 25:1904–1910.

Schilit Nitenson A, Stackpole EE, Truszkowski TL, Midroit M, Fallon JR, Bath KG (2015) Fragile X mental retardation protein regulates olfactory sensitivity but not odorant discrimination. Chemical senses 40:345–350.

Schoppa NE (2006) Synchronization of olfactory bulb mitral cells by precisely timed inhibitory inputs. Neuron 49:271–283.

Schoppa NE, Westbrook GL (2001) Glomerulus-specific synchronization of mitral cells in the olfactory bulb. Neuron 31:639–651.

Scotto-Lomassese S, Nissant A, Mota T, Neant-Fery M, Oostra BA, Greer CA, Lledo PM, Trembleau A, Caille I (2011) Fragile X mental retardation protein regulates new neuron differentiation in the adult olfactory bulb. The Journal of neuroscience: the official journal of the Society for Neuroscience 31:2205–2215.

Sinclair D, Oranje B, Razak KA, Siegel SJ, Schmid S (2017) Sensory processing in autism spectrum disorders and Fragile X syndrome-From the clinic to animal models. Neurosci Biobehav Rev 76:235–253.

Stopfer M, Bhagavan S, Smith BH, Laurent G (1997) Impaired odour discrimination on desynchronization of odour-encoding neural assemblies. Nature 390:70–74.

Takarae Y, Sweeney J (2017) Neural Hyperexcitability in Autism Spectrum Disorders. Brain Sci 7.

Telias M (2019) Molecular Mechanisms of Synaptic Dysregulation in Fragile X Syndrome and Autism Spectrum Disorders. Front Mol Neurosci 12:51.

Vaaga CE, Westbrook GL (2016) Parallel processing of afferent olfactory sensory information. The Journal of physiology 594:6715–6732.

Vislay RL, Martin BS, Olmos-Serrano JL, Kratovac S, Nelson DL, Corbin JG, Huntsman MM (2013) Homeostatic responses fail to correct defective amygdala inhibitory circuit maturation in fragile X syndrome. The Journal of neuroscience: the official journal of the Society for Neuroscience 33:7548–7558.

Wen TH, Lovelace JW, Ethell IM, Binder DK, Razak KA (2019) Developmental Changes in EEG Phenotypes in a Mouse Model of Fragile X Syndrome. Neuroscience 398:126–143.

Yang C et al. (2022) Restoration of FMRP expression in adult V1 neurons rescues visual deficits in a mouse model of fragile X syndrome. Protein Cell 13:203–219.

Yokoi M, Mori K, Nakanishi S (1995) Refinement of odor molecule tuning by dendrodendritic synaptic inhibition in the olfactory bulb. Proceedings of the National Academy of Sciences of the United States of America 92:3371–3375.

Zak JD, Schoppa NE (2022) Neurotransmitter regulation rather than cell-intrinsic properties shapes the high-pass filtering properties of olfactory bulb glomeruli. The Journal of physiology 600:393–417.

